# Primate phylogenomics uncovers multiple rapid radiations and ancient interspecific introgression

**DOI:** 10.1101/2020.04.15.043786

**Authors:** Dan Vanderpool, Bui Quang Minh, Robert Lanfear, Daniel Hughes, Shwetha Murali, R. Alan Harris, Muthuswamy Raveendran, Donna M. Muzny, Richard A. Gibbs, Kim C. Worley, Jeffrey Rogers, Matthew W. Hahn

## Abstract

Our understanding of the evolutionary history of primates is undergoing continual revision due to ongoing genome sequencing efforts. Bolstered by growing fossil evidence, these data have led to increased acceptance of once controversial hypotheses regarding phylogenetic relationships, hybridization and introgression, and the biogeographical history of primate groups. Among these findings is a pattern of recent introgression between species within all major primate groups examined to date, though little is known about introgression deeper in time. To address this and other phylogenetic questions, here we present new reference genome assemblies for three Old World Monkey species: *Colobus angolensis ssp. palliatus* (the black and white colobus), *Macaca nemestrina* (southern pig-tailed macaque), and *Mandrillus leucophaeus* (the drill). We combine these data with 23 additional primate genomes to estimate both the species tree and individual gene trees using thousands of loci. While our species tree is largely consistent with previous phylogenetic hypotheses, the gene trees reveal high levels of genealogical discordance associated with multiple primate radiations. We use strongly asymmetric patterns of gene tree discordance around specific branches to identify multiple instances of introgression between ancestral primate lineages. In addition, we exploit recent fossil evidence to perform fossil-calibrated molecular dating analyses across the tree. Taken together, our genome-wide data help to resolve multiple contentious sets of relationships among primates, while also providing insight into the biological processes and technical artifacts that led to the disagreements in the first place.

## Introduction

Understanding the history of individual genes and whole genomes is an important goal for evolutionary biology. It is only by understanding these histories that we can understand the origin and evolution of traits—whether morphological, behavioral, or biochemical. Until recently, our ability to address the history of genes and genomes was limited by the availability of comparative genomic data. However, genome sequences are now being generated extremely rapidly. In primates alone, there are already 23 species with published reference genome sequences and associated annotations (Table S1), as well as multiple species with population samples of whole genomes [1–11]. These data can now be used to address important evolutionary questions.

Several studies employing dozens of loci sampled across broad taxonomic groups have provided rough outlines of the evolutionary relationships and divergence times among primates [12,13]. Due to the rapid nature of several independent radiations within primates, these limited data cannot resolve species relationships within some clades [12–14]. For instance, the New World Monkeys (NWM) experienced a rapid period of diversification ~15-18 million years ago (mya) [15] (Figure 1), resulting in ambiguous relationships among the three Cebidae subfamilies (Cebinae=squirrel monkeys and capuchins, Aotinae=owl monkeys, and Callitrichinae=marmosets and tamarins) [12–14,16–18]. High levels of incomplete lineage sorting (ILS) driven by short times between the divergence of distinct lineages have led to a large amount of gene tree discordance in the NWM, with different loci favoring differing relationships among taxa. Given the known difficulties associated with resolving short internodes [19–21], as well as the multiple different approaches and datasets used in these analyses, the relationships among cebid subfamilies remain uncertain.

**Figure 1.**
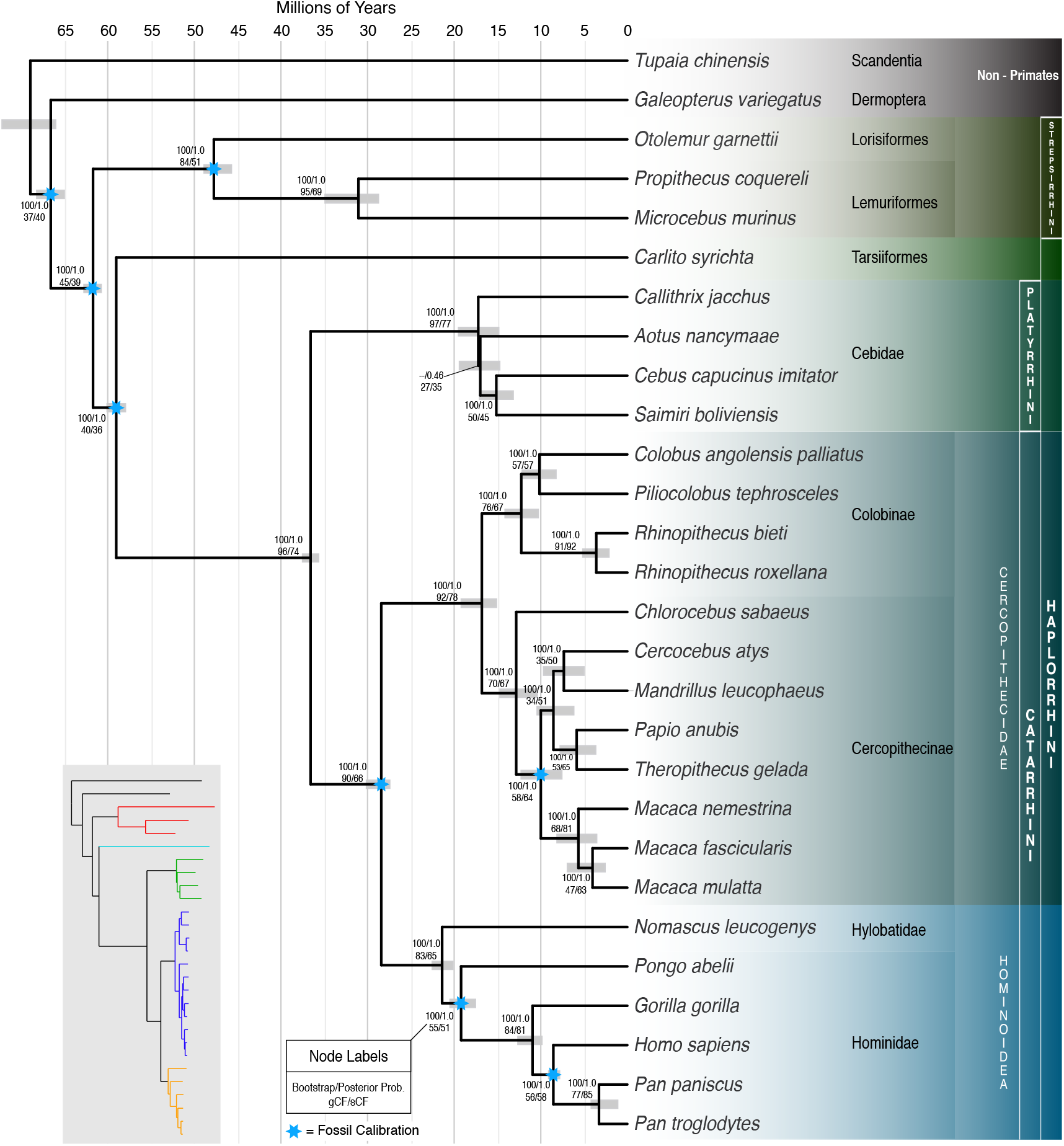
Species tree estimated using ASTRAL III with 1,730 gene trees (the *Mus musculus* outgroup was removed to allow for a visually finer scale). Common names for each species can be found in Table S1. Node labels indicate the bootstrap value from a maximum likelihood analysis of the concatenated dataset as well as the local posterior probability from the ASTRAL analysis. Gene concordance factors (gCF) and site concordance factors (sCF) are also reported. Eight fossil calibrations (blue stars; Table S5) were used to calibrate node ages. Grey bars indicate the minimum and maximum mean age from independent dating estimates. The inset tree with colored branches shows the maximum likelihood branch lengths estimated using a partitioned analysis of the concatenated alignment. Colors correspond to red = Strepsirrhini, cyan = Tarsiiformes, green = Platyrrhini (New World Monkeys), blue = Cercopithecoidea (Old World Monkeys), orange = Hominoidea (Apes).

In addition to issues of limited data and rapid radiations, a history of hybridization and subsequent gene flow between taxa means that there is no single dichotomously branching tree that all genes follow. Although it once was thought to be relatively rare (especially among animals, [22]), genomic studies have uncovered widespread patterns of recent introgression across the tree of life [23]. Evidence for recent or ongoing gene flow is especially common among the primates (e.g. [9,24–27]), sometimes with clear evidence for adaptive introgression (e.g. [28–30]). Whether widespread gene flow among primates is emblematic of their initial radiation (which began 60-75 mya, [13,31–33]) or is a consequence of current conditions—which include higher environmental occupancy and more secondary contact—remains an open question [34].

Here we report the sequencing and annotation of three new primate genomes, all Old World Monkey (OWM) species: *Colobus angolensis ssp. palliatus* (the black and white colobus), *Macaca nemestrina* (southern pig-tailed macaque), and *Mandrillus leucophaeus* (the drill). Together with the published whole genomes of extant primates, we present a phylogenomic analysis including 26 primate species and several closely related non-primates. Incorporating recently discovered fossil evidence [35], we perform fossil-calibrated molecular dating analyses to estimate divergence times, including dates for the crown primates as well as the timing of more recent splits. Compared to recent hybridization, introgression that occurred between two or more ancestral lineages (represented by internal branches on a phylogeny) is difficult to detect. To get around this limitation, we modify a previously proposed method for detecting introgression [36] and apply it to our whole-genome datasets, finding additional evidence for gene flow among ancestral primates. Finally, we closely examine the genealogical patterns left behind by the NWM radiation, as well as the biases of several methods that have been used to resolve this topology. We use multiple approaches to provide a strongly supported history of the NWM and primates in general, while also highlighting the large amounts of gene tree discordance across the tree caused by ILS and introgression.

## Results and Discussion

### Primate Genome Sequencing

The assembly and annotation of each of the three species sequenced for this project are summarized here, with further details listed in Table 1. A summary of all published genomes used in this study, including links to the assemblies and NCBI BioProjects, is available in Table S2.

**Table 1.** Genomes sequenced in this study and associated assembly and annotation metrics. BUSCO percentages reflect the complete and fragmented genes relative to the Euarchontoglires ortholog database v9.

| Species name | Assembly Accession | Assembly Total length | Number of scaffolds | Scaffold N50 | Number of contigs | Protein-coding genes | BUSCO |
| --- | --- | --- | --- | --- | --- | --- | --- |
| *Colobus angolensis ssp. palliatus* (the black and white colobus) | GCF_000951035.1 | 2,970,124,662 | 13,124 | 7,840,981 | 197,124 | 20,222 | 95.82% |
| *Macaca nemestrina* (pig-tailed macaque) | GCF_000956065.1 | 2,948,703,511 | 9,733 | 15,219,753 | 94,057 | 21,017 | 95.98% |
| *Mandrillus leucophaeus* (drill) | GCF_000951045.1 | 3,061,992,840 | 12,821 | 3,186,748 | 246,054 | 20,465 | 95.45% |

The sequencing effort for *Colobus angolensis ssp. palliatus* produced 514 Gb of data, which are available in the NCBI Short Read Archive (SRA) under the accession SRP050426 (BioProject PRJNA251421). Assembly of these data resulted in a total assembly length of 2.97 Gb in 13,124 scaffolds (NCBI assembly Cang.pa_1.0; GenBank accession GCA_000951035.1) with an average per base coverage of 86.8X. Subsequent annotation via the NCBI Eukaryotic Genome Annotation Pipeline (annotation release ID: 100) resulted in the identification of 20,222 protein-coding genes and 2,244 non-coding genes. An assessment of the annotation performed using BUSCO 3.0.2 [37] in conjunction with the Euarchontoglires ortholog database 9 (https://busco-archive.ezlab.org/v3/datasets/euarchontoglires_odb9.tar.gz) indicated that 95.82% complete or fragmented single-copy orthologs (91.68% complete, 4.13% fragmented) were present among the annotated protein-coding genes. Comprehensive annotation statistics for *C. angolensis ssp. palliatus* with links to the relevant annotation products available for download can be viewed at https://www.ncbi.nlm.nih.gov/genome/annotation_euk/Colobus_angolensis_palliatus/100/.

For *Macaca nemistrina*, 1,271 Gb of data were produced (SRA accession SRP045960; BioProject PRJNA2791) resulting in an assembled genome length of 2.95 Gb in 9,733 scaffolds (Mnem_1.0; GenBank accession GCF_000956065.1). This corresponds to an average per base coverage of 113.1X when both short and long-read data are combined (Materials and Methods). The NCBI annotation resulted in 21,017 protein coding genes and 13,163 non-coding genes (annotation release ID: 101). A BUSCO run to assess the completeness of the annotation (as above) indicated 95.98% complete or fragmented single-copy orthologs (92.23% complete, 3.75% fragmented) present among the annotated protein-coding genes. Comprehensive annotation statistics for *M. nemistrina* with links to the relevant annotation products available for download can be viewed at https://www.ncbi.nlm.nih.gov/genome/annotation_euk/Macaca_nemestrina/101/.

Sequencing of *Mandrillus leucophaeus* libraries resulted in 334.1 Gb of data (SRA accession SRP050495; BioProject PRJNA251423) that once assembled resulted in a total assembly length of 3.06 Gb in 12,821 scaffolds (Mleu.le_1.0; GenBank accession GCA_000951045.1) with an average coverage of 117.2X per base. The NCBI annotation produced of 20,465 protein coding genes and 2,300 non-coding genes (annotation release ID: 100). A BUSCO run to assess the completeness of the annotation (as above) indicated 95.45% complete or fragmented single-copy orthologs (91.38% complete, 4.07% fragmented) present among the annotated protein-coding genes. The full annotation statistics with links to the associated data can be viewed at https://www.ncbi.nlm.nih.gov/genome/annotation_euk/Mandrillus_leucophaeus/100/.

### Phylogenetic Relationships Among Primates

To investigate phylogenetic relationships among primates, we selected the longest isoform for each protein-coding gene from 26 primate species and 3 non-primate species (Table S1). After clustering, aligning, trimming, and filtering (Materials and Methods) there were 1,730 single-copy orthologs present in at least 27 of the 29 species. These cutoffs ensure high species coverage while still retaining a large number of orthologs. The coding sequences of these orthologs have an average length of 1,018 bp and 178 parsimony-informative characters per gene. Concatenation of these loci resulted in an alignment of 1,761,114 bp, with the fraction of gaps/ambiguities varying from 4.04% (*Macaca mulatta*) to 18.37% (*Carlito syrichta*) (Table S3). We then inferred the species tree using both gene tree (as implemented in ASTRAL III, [38] and concatenation (as implemented in IQ-TREE 2; [39]) approaches.

We inferred 1,730 individual gene trees from nucleotide alignments using maximum likelihood in IQ-TREE 2, and then used these gene tree topologies as input to ASTRAL III (Materials and Methods). We used the mouse, *Mus musculus*, as an outgroup to root the species tree. This approach resulted in a topology (which we refer to as “ML-ASTRAL”; Figure 1) that largely agrees with previously published phylogenies [12,13]. Maximum likelihood analysis of the concatenated nucleotide alignment (which we refer to as “ML-CONCAT”) using IQ-TREE resulted in a topology that differed from the ML-ASTRAL tree only with respect to the placement of *Aotus nancymaae* (owl monkey): rather than sister to the *Saimiri+Cebus* clade (as in Figure 1), the ML-CONCAT tree places *Aotus* sister to *Callithrix jacchus*, a minor rearrangement around a very short internal branch (Figure 1). All branches of the ML-ASTRAL species tree are supported by maximum local posteriors [40], except for the branch that defines *Aotus* as sister to the *Saimiri+Cebus* clade (0.46 local posterior probability). Likewise, each branch in the ML-CONCAT tree is supported by 100% bootstrap values, including the branch uniting *Aotus* and *Callithrix*. We return to this conflict in the next section.

There has been some contention as to the placement of the mammalian orders Scandentia (treeshrews) and Dermoptera (colugos) [41–50]. Both the ML-ASTRAL and ML-CONCAT trees place these two groups outside the Primates with maximal statistical support (i.e. local posterior probabilities of 1.0 and bootstrap values of 100%; Figure 1), with Dermoptera as the closest sister lineage to the Primates [12,51–53]. However, while support values such as the bootstrap or posterior provide statistical confidence in the species tree topology, there can be large amounts of underlying gene tree discordance even for branches with 100% support (e.g. [54–56]). To assess discordance generally, and the relationships among the Primates, Scandentia, and Dermoptera in particular, we used IQ-TREE to calculate both gene (gCF) and site (sCF) concordance factors [57] for each internal branch of the topology in Figure 1. These two measures represent the fraction of genes and sites, respectively, that are in agreement with the species tree for any particular branch.

Examining concordance factors helps to explain previous uncertainty in the relationships among Primates, Scandentia, and Dermoptera (Figure 1). Although the bootstrap support is 100% and the posterior probability is 1.0 on the branch leading to the Primate common ancestor, the gene concordance factor is 45% and the site concordance factor is 39%. These values indicate that, of decisive gene trees (*n*=1663), only 45% of them contain the branch that is in the species tree; this branch reflects the Primates as a single clade that excludes Scandentia and Dermoptera. While the species tree represents the single topology supported by the most gene trees (hence the strong statistical support for this branch), the concordance factors also indicate that a majority of individual topologies have histories that differ from the estimated species tree. In fact, the gCF value indicates that 55% of trees do not support a monophyletic Primate order, with either Dermoptera, Scandentia, or both lineages placed within Primates. Likewise, the sCF value indicates that only 39% of parsimony-informative sites in the total alignment support the branch uniting all primates, with 30% favoring Dermoptera as sister to the Primate sub-order Strepsirrhini and 31% placing Dermoptera sister to the Primate sub-order Haplorrhini. Similarly, only a small plurality of genes and sites have histories that place Dermoptera as sister to the Primates rather than either of the two alternative topologies (gCF=37, sCF=40; Figure 1), despite the maximal statistical support for these relationships. While discordance at individual gene trees can result from technical problems in tree inference (e.g. long-branch attraction, low phylogenetic signal, poorly aligned sequences, or model misspecification), it also often reflects biological causes of discordance such as incomplete lineage sorting and introgression. The fraction of discordant gene trees for branches near the base of the primate tree is no larger than the fraction on branches reflecting more recent radiations (Figure 1), and therefore likely results from both technical errors and the biological consequences of the rapid radiation of mammalian lineages during this period [32,51,58,59].

Within the Primates, the phylogenetic affiliation of tarsiers (represented here by *Carlito syrichta*) has been debated since the first attempts by Buffon (1765) and Linnaeus (1767-1770) to systematically organize described species [60]. Two prevailing hypotheses group tarsiers (Tarsiiformes) with either lemurs and lorises (the “prosimian” hypothesis, [61]) or with Simiiformes (the “Haplorrhini” hypothesis, [62], where Simiiformes = Apes+OWM+NWM). The ML-ASTRAL and ML-CONCAT analyses place Tarsiiformes with Simiiformes, supporting the Haplorrhini hypothesis (Figure 1). The strepsirrhines come out as a well-supported group sister to the other primates. Again, our inference of species relationships is consistent with previous genomic analyses [58,63], but also highlights the high degree of discordance in this part of the tree. The rapid radiation of mammalian lineages that occurred in the late Paleocene and early Eocene [32] encompassed many of the basal primate branches, including the lineage leading to Haplorrhini. The complexity of this radiation is likely the reason for low gCF and sCFs (39.5% and 36%, respectively) for the branch leading to Haplorrhini, and perhaps explains why previous studies recovered conflicting resolutions for the placement of tarsiers [31,64,65].

The remaining branches of the species tree that define major primate clades all have remarkably high concordance with the underlying gene trees (gCF > 80%), though individual branches within these clades do not. The gCFs for the branches defining these clades are: Strepsirrhini (lemurs+lorises) = 84.5, Catarrhini (OWM+Apes) = 90.0, Platyrrhini (NWM) = 96.6, Hominoidea (Apes) = 82.7, and Cercopithecidae (OWM) = 92.3 (Figure 1). High gene tree/species tree concordance for these branches is likely due to a combination of more recent divergences (increasing gene tree accuracy) and longer times between branching events [66]. Within these clades, however, we see multiple recent radiations. One of the most contentious has been among the New World Monkeys, a set of relationships we address next.

### Concatenation Affects Resolution of the New World Monkey Radiation

Sometime during the mid-to-late Eocene (~45-34 mya), a small number of primates arrived on the shores of South America [15,67]. These monkeys likely migrated from Africa [67] and on arrival underwent multiple rounds of extinction and diversification [15], the most recent of which was aided by a period of warming referred to as the Mid-Miocene Climatic Optimum [59]. Three extant families from this radiation now make up the New World Monkeys (Platyrrhini, Figure 1). Because of the rapidity with which these species spread and diversified across the new continent, relationships at the base of the NWM have been hard to determine [12–14,16–18].

As reported above, the concatenated analysis (ML-CONCAT) gives a different topology than the gene tree-based analysis (ML-ASTRAL). Specifically, the ML-CONCAT analysis supports a symmetrical tree, with *Aotus* sister to *Callithrix* (Figure 2A). In contrast, ML-ASTRAL supports an asymmetrical (or “caterpillar”) tree, with *Aotus* sister to a clade comprised of *Saimiri+Cebus* (Figure 2B). There are reasons to have doubts about both topologies. It is well known that carrying out maximum likelihood analyses of concatenated datasets can result in incorrect species trees, especially when the time between speciation events is short [68,69]. In fact, the specific error that is made in these cases is for ML concatenation methods to prefer a symmetrical four-taxon tree over an asymmetrical one, exactly as is observed here. Gene tree-based methods such as ASTRAL are not prone to this particular error, as long as the underlying gene trees are all themselves accurate [70,71]. However, if there is bias in gene tree reconstruction, then there are no guarantees as to the accuracy of the species tree. In addition, the ML-ASTRAL tree is supported by only a very small plurality of gene trees: there are 442 trees supporting this topology, compared to 437 supporting the ML-CONCAT topology and 413 supporting the third topology (Figure 2D). This small excess of supporting gene trees also explains the relatively low posterior support for this branch in the species tree (Figure 1). Additionally, a polytomy test [72], implemented in ASTRAL and performed using ML gene trees, failed to reject the null hypothesis of “polytomy” for the branch uniting *Aotus+*(*Saimiri,Cebus*) (*P*=0.47).

**Figure 2.**
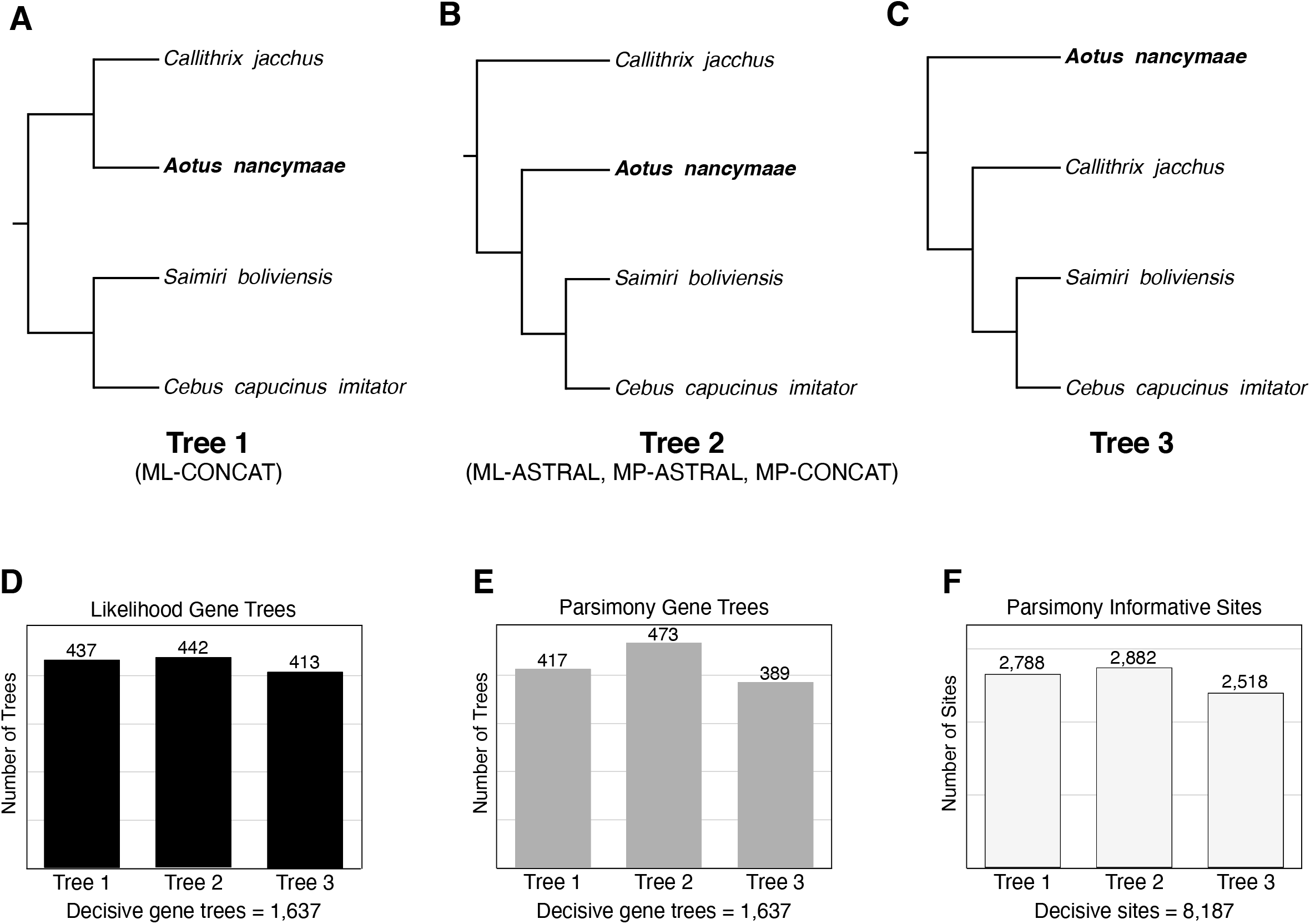
The three most frequent topologies of New World Monkeys. A) Tree 1 is the symmetrical topology inferred by the maximum likelihood concatenated analysis (ML-CONCAT) of 1,730 loci (1.76 Mb). B) Tree 2 is the asymmetrical topology inferred by ASTRAL III using either maximum likelihood (ML-ASTRAL) or maximum parsimony (MP-ASTRAL) gene tree topologies. Using maximum parsimony on the concatenated alignment also returns this tree (MP-CONCAT). C) Tree 3 is the alternative resolution recovered at high frequency in all gene tree analyses, though it is not the optimal species tree using any of the methods. D) Number of gene trees supporting each of the three resolutions of the NWM clade when maximum likelihood is used to infer gene tree topologies. There are 1,637 decisive gene trees for these splits. E) Gene tree counts when maximum parsimony is used to infer gene tree topologies. F) Number of parsimony informative sites in the concatenated alignment supporting each of the three resolutions.

To investigate these relationships further, we carried out additional analyses. The trees produced from concatenated alignments are biased when maximum likelihood is used for inference, but this bias does not affect parsimony methods [21,73]. Therefore, we analyzed exactly the same concatenated 1.76 Mb alignment used as input for ML, but carried out a maximum parsimony analysis in PAUP* [74]. As would be expected given the known biases of ML methods, the maximum parsimony tree (which we refer to as “MP-CONCAT”) returns the same tree as ML-ASTRAL, supporting an asymmetric topology of NWMs (Figure 2B). Underlying this result is a relatively large excess of parsimony-informative sites supporting this tree (Figure 2F), which results in maximal bootstrap values for every branch. The two most diverged species in this clade (*Saimiri* and *Callithrix*) are only 3.26% different at the nucleotide level, so there should be little effect of multiple substitutions on the parsimony analysis.

As mentioned above, gene tree-based methods (such as ASTRAL) are not biased when accurate gene trees are used as input. However, in our initial analyses we used maximum likelihood to infer the individual gene trees. Because protein-coding genes are themselves often a combination of multiple different underlying topologies [75], ML gene trees may be biased, and using them as input to gene tree-based methods may still lead to incorrect inferences of the species tree [76]. Therefore, we used the same 1,730 loci as above to infer gene trees using maximum parsimony with MPBoot [77]. Although the resulting topologies still possibly represent the average over multiple topologies contained within a protein-coding gene, using parsimony ensures that this average tree is not a biased topology. These gene trees were used as input to estimate a species tree using ASTRAL; we refer to this as the “MP-ASTRAL” tree. Once again, the methods that avoid known biases of ML lend further support to an asymmetric tree, placing *Aotus* sister to the *Saimiri+Cebus* clade (Figure 2B). In fact, the gene trees inferred with parsimony now show a much greater preference for this topology, with a clear plurality of gene trees supporting the species tree (473 vs. 417 supporting the second-most common tree; Figure 2E). As a consequence, the local posterior for this branch in the MP-ASTRAL tree is 0.92 and the polytomy test performed using MP gene trees rejects (*p* = 0.037) the null hypothesis of “polytomy” for the branch uniting *Aotus+*(*Saimiri,Cebus*). The increased number of concordant gene trees using parsimony suggests that the gene trees inferred using ML may well have been suffering from the biases of concatenation within each locus, reducing the observed levels of concordance.

A recent analysis of NWM genomes found *Aotus* sister to *Callithrix*, as in the ML-CONCAT tree, despite the use of gene trees to build the species tree [18]. However, the outgroup used in this analysis is a closely related species (*Brachyteles arachnoides*) that diverged during the NWM radiation and that shares a recent common ancestor with the ingroup taxa [12,13]. If the outgroup taxon used to root a tree shares a more recent common ancestor with subsets of ingroup taxa at an appreciable number of loci, the resulting tree topologies will be biased. A similar problem likely arose in previous studies that have used the Scandentia or Dermoptera as outgroups to Primates. In general, this issue highlights the difficulty in choosing outgroups: though we may have 100% confidence that a lineage lies outside our group of interest in the species tree, a reliable outgroup must also not have any discordant gene trees that place it inside the ingroup.

### Strongly Supported Divergence Times Using Fossil Calibrations

Fossil-constrained molecular dating was performed using 10 independent datasets, each of which consisted of 40 protein-coding genes randomly selected (without replacement) and concatenated. The resulting datasets had an average alignment length of 39,374 bp (SD=2.6×10^3^, Table S4). Although individual discordant trees included in this analysis may have different divergence times, the difference in estimates of dates should be quite small [78]. We used eight dated fossils (blue stars in Figure 1) from 10 studies for calibration (Table S5). The most recent of these fossils is ~5.7 mya [79], while the most ancient is 55.8 mya [80]. Each separate dataset and the same set of “soft” fossil constraints, along with the species tree in Figure 1, were used as input to PhyloBayes 3.3 [81] which was run twice to assess convergence (Materials and Methods).

We observed tight clustering of all estimated node ages across datasets and independent runs of PhyloBayes (Figure 3 and Table S6). In addition, the ages of most major crown nodes estimated here are largely in agreement with previously published age estimates (Table 2). Some exceptions include the age of the crown Strepsirrhini (47.4 mya) and Haplorrhini (59.0 mya) which are more recent than many previous estimates for these nodes (range in the literature is Strepsirrhini = 51.6 - 68.7, Haplorrhini = 60.6 – 81.3). The crown nodes for Catarrhini, Hominoidea, and Cercopithecidae (28.4, 21.4, and 16.8 mya, respectively) all fall within the range of variation recovered in previous studies (Table 2).

**Figure 3.**
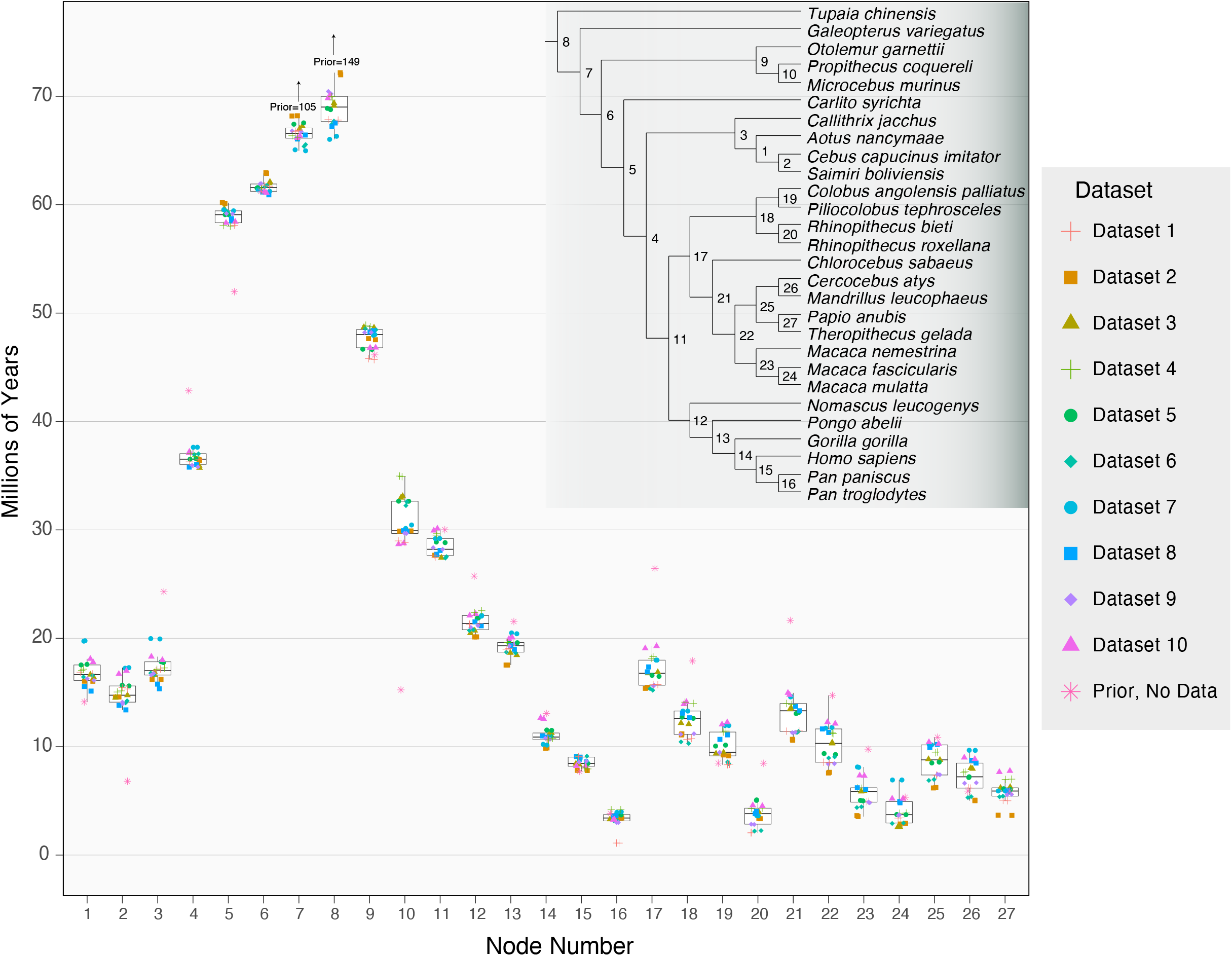
Mean node ages for independent Phylobayes dating runs on 10 different datasets (each dataset was run twice). Box plots show the median, interquartile range, and both minimum and maximum values of the mean nodes ages. An additional run was performed with no sequence data to ascertain the prior on node divergence times in the presence of fossil calibrations (pink asterisks). Some prior ages were too large to include in the plot while still maintaining detail; these ages are given as numeric values. The species tree topology is from Figure 1, 95% highest posterior density (HPD) intervals for each node are reported in Table S6.

**Table 2.**

| Node | This Study | This Study No Max* | Herrera *et al.* [32] | Kistler *et al.* [33] | Perez *et al.* [17] | Springer *et al.* [13] | Meredith *et al.* [45] | Perelman *et al.* [12] | Wilkinson *et al.* [83] | Chatterjee *et al.* [31] |
| --- | --- | --- | --- | --- | --- | --- | --- | --- | --- | --- |
| Primates | 61.7 | 67.5 | 63.9 | 68 | NA | 67.8 | 71.5 | 87.2 | 84.5 | 63.7 |
| Strepsirrhini | 47.4 | 50.2 | 61.4 | 59 | NA | 54.2 | 55.1 | 68.7 | 49.8 | 51.6 |
| Haplorrhini | 59.0 | 63.8 | 61.9 | 67 | 60.6 | 61.2 | 62.4 | 81.3 | NA | NA |
| Catarrhini | 28.4 | 29.0 | 32.1 | 33 | 27.8 | 25.1 | 20.6 | 31.6 | 31.0 | 29.3 |
| Hominoidea | 21.4 | 21.6 | NA | 21 | 18.44 | 17.4 | 14.4 | 20.3 | NA | 21.5 |
| Cercopithecidae | 16.8 | 16.9 | NA | 24 | 13.4 | 13.2 | NA | 17.6 | 14.1 | 23.4 |

Mean crown node divergence times estimated in this study compared with mean divergences times estimated by eight prior studies. Estimates were calculated by averaging the mean times across all runs for 10 independent datasets. \*Refers to the average divergence time of the crown node for the indicated taxonomic group when the 65.8 my maximum constraint was removed from the Primate node.

Our estimate for the most recent common ancestor of the extant primates (i.e. the last common ancestor of Haplorrhini and Strepsirrhini) is 61.7 mya, which is slightly more recent than several studies [13,31,33,82] and much more recent than other studies [12,83,84]. However, our estimate is in good agreement with Herrera *et al*. [32], who used 34 fossils representing extinct and extant lineages (primarily Strepsirrhines) to infer divergence times among primates, concluding that the split occurred approximately 64 mya. One similarity between our study and that of Herrera *et al*. is that we have both used the maximum constraint of 65.8 my on the ancestral primate node suggested by Benton *et al*. [85], which likely contributes to the more recent divergence. It is worth noting that the soft bounds imposed in our analysis permit older ages to be sampled from the Markov chain, but these represented only a small fraction (median 3.37%) of the total sampled states after burn-in (Table S5). To determine the effects of imposing the 65.8 my maximum constraint on the Primate node, we analyzed all 10 datasets for a third time with this constraint removed and report the divergence time of major primate clades in Table 2 (“No Max” entries).

There are several caveats to our age estimates that should be mentioned. Maximum age estimates for the crown node of any given clade are defined by the oldest divergence among sampled taxa in the clade. This limitation results in underestimates for nearly all crown node ages as, in practice, complete taxon sampling is difficult to achieve. Fossil calibrations are often employed as minimum constraints in order to overcome the limitations imposed by taxon sampling, allowing older dates to be estimated more easily. On the other hand, the systematic underestimation of crown node ages due to taxon sampling is somewhat counteracted by the overestimation of speciation times due to ancestral polymorphism. Divergence times estimated from sequence data represent the coalescence times of sequences, which are necessarily older than the time at which two incipient lineages diverged [86,87]. This overestimation will have a proportionally larger effect on recent nodes (such as the *Homo/Pan* split; Figure 3, node 15), but the magnitude can be no larger than the average level of polymorphism in ancestral populations and will be additionally reduced by post-divergence gene flow.

### Introgression During the Radiation of Primates

There is now evidence for recent inter-specific gene flow between many extant primates, including introgression events involving humans [24], gibbons [88,89], baboons [9,27], macaques [90,91], and vervet monkeys [10], among others. While there are several widely used methods for detecting introgression between closely related species (see chapters 5 and 9 in [92]), detecting ancient gene flow is more difficult. One of the most popular methods for detecting recent introgression is the *D* test (also known as the “ABBA-BABA” test; [93]). This test is based on the expectation that, for any given branch in a species tree, the two most frequent alternative resolutions should be present in equal proportions. However, the *D* test uses individual SNPs to evaluate support for alternative topologies, and explicitly assumes an infinite sites model of mutation (i.e. no multiple hits). As this assumption will obviously not hold the further back in time one goes, a different approach is needed.

Fortunately, Huson *et al*. [36] described a method that uses gene trees themselves (rather than SNPs) to detect introgression. Using the same expectations as in the *D* test, these authors looked for a deviation from the expected equal numbers of alternative tree topologies using a test statistic they refer to as Δ. As far as we are aware, Δ has only rarely been used to test for introgression in empirical data, possibly because of the large number of gene trees needed to assess significance, or the assumptions of the parametric method proposed to obtain *P*-values. Here, given our large number of gene trees and large number of internal branches to be tested, we adapt the Δ test for genome-scale data.

To investigate patterns of introgression within primates, we used 1,730 single-copy loci to test for deviations from the null expectation of Δ on each of the 24 internal branches of the primate phylogeny (Materials and Methods). To test whether deviations in Δ were significant (i.e. Δ > 0), we generated 2000 resampled datasets of 1,730 gene tree topologies each. *P*-values were calculated from *Z*-scores generated from these resampled datasets. Among the 17 branches where at least 5% of topologies were discordant, we found 7 for which Δ had *P*<0.05.

To further verify these instances of potential introgression, for each of these seven branches we increased the number of gene trees used by subsampling a smaller set of taxa. We randomly chose four taxa for each internal branch tested that also had this branch as an internal branch, and then re-aligned orthologs present in a single copy in each taxon. These steps resulted in ~3,600-6,400 genes depending on the branch being tested (Supplementary Table S7). Additionally, because instances of hybridization and introgression are well documented among macaques [90,91,94], we similarly re-sampled orthologs from the three *Macaca* species in our study.

We recalculated Δ using the larger gene sets and found significant evidence (after correcting for *m*=17 multiple comparisons by using a cutoff of *P* = 0.00301) for six introgression events, all of which occurred among the Papionini (Figure 4 and see next paragraph). Within the Hominoidea, we found Δ = 0.0518 for the branch leading to the great apes, but *P* = 0.030. The asymmetry in gene tree topologies here suggests gene flow may have happened between gibbons (represented by *Nomascus*) and the ancestral branch leading to the African hominoids (humans, chimpanzees, and gorillas), but, like the *D* test, Δ cannot tell us the direction of introgression. Although currently separated by significant geographic distances (African apes south of the Sahara Desert and gibbons all in southeast Asia), it is worth noting that fossil hominoids dating from the early to late Miocene had a broad distribution extending from southern Africa to Europe and Asia [95]. Support for introgression between ancestral hominins and ancestral chimpanzees has been previously reported [96]; our four-taxon analyses found marginal support for this conclusion (Δ = 0.0917, *P* = 0.055).

**Figure 4.**
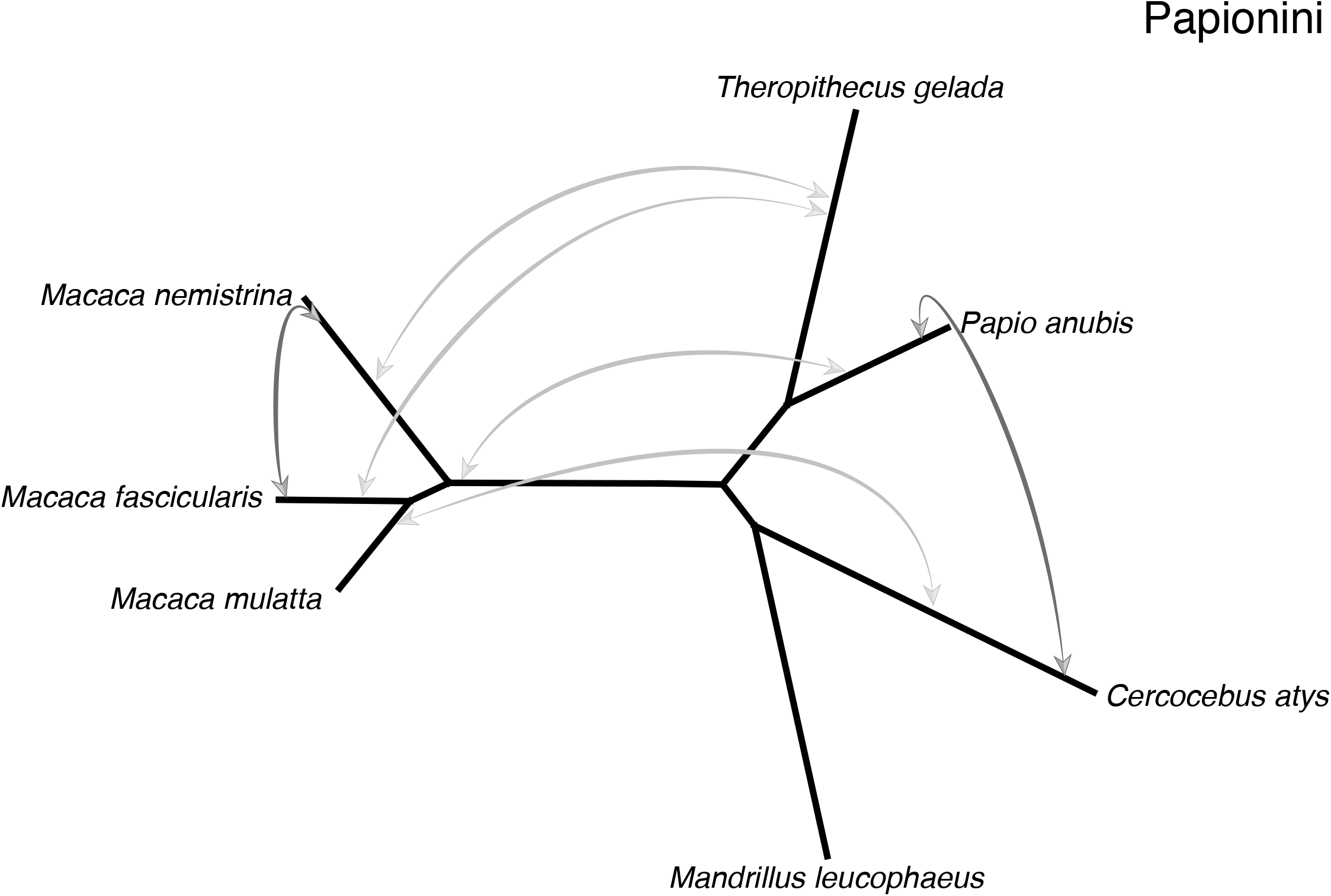
Introgression among Papionini taxa (the species tree is unrooted for clarity). Arrows indicate that a significant Δ was found in our four taxon tests and identify the two lineages inferred to have exchanged genes (values underlying these tests are listed in Table S7). Among the Papionini, there was evidence of introgression between African taxa (*Papio, Theropithecus*, and *Cercocebus*) and Asian *Macaca* species (light grey arrows). Introgression events likely occurred between African taxa and the ancestral *Macaca*, which had a wide distribution across northern Africa prior to the radiation throughout Asia 2-3 mya [131]. More recent instances of introgression are inferred between macaque species and among the African Papionini (dark grey arrows).

Within the OWM, ~40% of Cercopithicine species are known to hybridize in nature [34]. Consistent with this, *Macaca nemestrina* and *M. fascicularis* showed a strong signature of gene flow in our data (Δ = 0.1761, *P* = 1.377e-09). These two species have ranges that currently overlap (Figure S1). In contrast to the clear signal of recent gene flow in the macaques, we detected a complex pattern of ancient introgression between the African Papionini (*Cercocebus, Mandrillus, Papio, and Theropithecus*) and the Asian Papionini (*Macaca*) (Figure 4). The Δ test was significant using multiple different subsamples of four taxa, suggesting multiple ancestral introgression events. An initial attempt to disentangle these events using Phylonet v3.8.0 [97,98] with the seven Papionini species and an outgroup was unsuccessful, as Phylonet failed to converge on an optimal network for these taxa. When there are multiple episodes of gene flow within a clade, even complex computational machinery may be unable to infer the correct combination of events.

As an alternative approach, we used four-taxon trees to estimate Δ for each *Macaca* species paired with two African Papionini (one from the *Papio+Theropithecus* clade and one from the *Mandrillus +Cercocebus* clade; see Table S7) and an outgroup. Significant introgression was detected using each of the *Macaca* species and three of the four African Papionini species (*Cercocebus, Theropithecus*, and *Papio*). These results suggest gene flow between the ancestor of the three *Macaca* species in our analysis and the ancestors of the three African Papionini in our analysis, or one introgression event involving the ancestor of all four African species coupled with a second event that masked this signal in *Mandrillus*. This second event may either have been biological (additional introgression events masking the signal), or technical (possibly the lack of continuity or completeness of the *Mandrillus* reference genome sequence), but in either case we could not detect introgression in the available drill sequence. The latter scenario would fit better with the current geographic distributions of these species, as they are on two different continents. However, the fossil record indicates that by the late Miocene to late Pleistocene the ancestral distribution of the genus *Macaca* covered all of North Africa, into the Levant, and as far north as the U.K. (Figure S1; [99]). The fossil record for *Theropithecus* indicates several species had distributions that overlapped with *Macaca* during this time, including in Europe and as far east as India (Figure S1, [100,101]). Ancestral macaques and ancestral papionins may therefore have come into contact in the area of the Mediterranean Sea. The Sahara Desert is also responsible for the current disjunct distributions of many of these species. However, this region has experienced periods of increased rainfall or “greenings” over the past several million years [102–104]. Faunal migration through the Sahara, including by hominins, is hypothesized to have occurred during these green periods [103,105,106] resulting in successive cycles of range expansion and contraction [107]. Hybridization and introgression could have occurred between the ancestors of these groups during one of these periods.

Our results on introgression come with multiple caveats, both about the events we detected and the events we did not detect. As with the *D* test, there are multiple alternative explanations for a significant value of Δ besides introgression. Ancestral population structure can lead to an asymmetry in gene tree topologies [108] though it requires a highly specific, possibly unlikely population structure. For instance, if the ancestral population leading to *Macaca nemestrina* was more closely related to *M. fascicularis* than was the ancestral population leading to its sister species, *M. mulatta* (Figure 4), then there could be an unequal number of alternative topologies. Similarly, any bias in gene tree reconstruction that favors one alternative topology over the other could potentially lead to a significant value of Δ. While this scenario is unlikely to affect recent divergences using SNPs, well known biases that affect topology reconstruction deeper in the tree (such as long-branch attraction) could lead to gene tree asymmetries. However, we did not observe any significant Δ-values for branches more than ~10 my old.

There are also multiple reasons why our approach may have missed introgression events, especially deeper in the tree. All methods that use asymmetries in gene tree topologies miss gene flow between sister lineages, as such events do not lead to changes in the proportions of underlying topologies. Similarly, equal levels of gene flow between two pairs of non-sister lineages can mask both events, while even unequal levels will lead one to miss the less-frequent exchange. More insidiously, especially for events further back in time, extinction of the descendants of hybridizing lineages will make it harder to detect introgression. Internal branches closer to the root will be on average longer than those near the tips because of extinction [109], and therefore introgression between non-sister lineages would have to occur longer after speciation in order to be detected. For instance, gene flow among Strepsirrhine species has been detected in many previous analyses of more closely related species (e.g. [110–113]) but the deeper relationships among the taxa sampled here may have made it very difficult to detect introgression. Nevertheless, our analyses were able to detect introgression between many primate species across the phylogeny.

## Conclusions

Several previous phylogenetic studies of primates have included hundreds of taxa, but fewer than 70 loci [12,13]. While the species tree topologies produced by these studies are nearly identical to the one recovered in our analysis, the limited number of loci meant that it was difficult to assess gene tree discordance accurately. By estimating gene trees from 1,700 single-copy loci, we were able to assess the levels of discordance present at each branch in the primate phylogeny. Understanding discordance helps to explain why there have been longstanding ambiguities about species relationships near the base of primates and in the radiation of New World Monkeys. Our analyses reveal how concatenation of genes—or even of exons—can mislead maximum likelihood phylogenetic inference in the presence of discordance, but also how to overcome the biases introduced by concatenation in some cases. Discordance also provides a window into introgression among lineages, and here we have found evidence for exchange among several species pairs. Each instance of introgression inferred from the genealogical data is plausible insofar as it can be reconciled with current and ancestral species distributions.

## Materials and Methods

### Source Material and Sequencing

For the sequencing of the *Colobus angolensis palliatus* genome, paired-end (100 bp) libraries were prepared using DNA extracted from heart tissue (isolate OR3802) kindly provided by Dr. Oliver Ryder (San Diego Zoo). Sequencing was performed using nine Illumina Hi-seq 2000 lanes and four Illumina Hi-seq 2500 lanes with subsequent assembly carried out using ALLPATHS-LG software (v. 48744) [114]. Additional scaffolding and gap-filling was performed using Atlas-Link v. 1.1 (https://www.hgsc.bcm.edu/software/atlas-link) and Atlas-GapFill v. 2.2. (https://www.hgsc.bcm.edu/software/atlas-gapfill) respectively. Annotation for all three species was carried out using the NCBI Eukaryotic Genome Annotation Pipeline. A complete description of the pipeline can be viewed at https://www.ncbi.nlm.nih.gov/genome/annotation_euk/process/.

For the sequencing of the *Macaca nemestrina* genome, DNA was extracted from a blood sample (isolate M95218) kindly provided by Dr. Betsy Ferguson and Dr. James Ha (Washington National Primate Research Center). Paired-end libraries were prepared and sequenced on 20 Illumina Hi-Seq 2000 lanes with the initial assembly performed using ALLPATHS-LG as above. Scaffolding was conducted using Atlas-Link v. 1.1. Additional gap-filling was performed using the original Illumina reads and Atlas-GapFill v. 2.2, as well as long reads generated using the Pacific Biosciences RS (60 SMRT cells) and RSII (50 SMRT cells) platforms. The PacBio reads were mapped to scaffolds to fill remaining gaps in the assembly using PBJelly2 (v. 14.9.9) [115].

For the sequencing of the *Mandrillus leucophaeus* genome, DNA was extracted from heart tissue (isolate KB7577) kindly provided by Dr. Oliver Ryder (San Diego Zoo). Paired-end libraries were prepared and sequenced on nine Illumina Hi-Seq 2000 lanes with the initial assembly performed using ALLPATHS-LG as above. Additional scaffolding was completed using Atlas-Link v. 1.1 and additional gap-filling in scaffolds was performed using the original Illumina reads and Atlas-GapFill v. 2.2.

### Phylogenomic Analyses

The full set of protein-coding genes for 26 primates and 3 non-primates were obtained by combining our newly sequenced genomes with already published data (see Table S1 for references and accessions and Tables 1 and S2 for genome statistics). Ortholog clustering was performed by first executing an all-by-all BLASTP search [116,117] using the longest isoform of each protein coding gene from each species. The resulting BLASTP output was clustered using the mcl algorithm [118] as implemented in FastOrtho [119] with various inflation parameters (the maximum number of clusters was obtained with *inflation=5*). Orthogroups were then parsed to retain those genes present as a single-copy in all 29 taxa (1,180 genes), 28 of 29 taxa (1,558 genes), and 27 of 29 taxa (1,735 genes). We chose to allow up to two missing species per alignment to maximize the data used in our phylogenomic reconstructions while maintaining high taxon-occupancy in each alignment. Coding sequences (CDS) for each single-copy orthogroup were aligned, cleaned, and trimmed via a multi-step process: First, sequences in each orthogroup were aligned by codon using GUIDANCE2 [120] in conjunction with MAFFT v7.407 [121] with 60 bootstrap replicates. Sequence residues in the resulting MAFFT alignment with GUIDANCE scores < 0.93 were converted to gaps and sites with > 50% gaps were removed using Trimal v1.4.rev22 [122]. Alignments shorter than 200 bp (full dataset) or 300 bp (four-taxon tests for introgression), and alignments that were invariant or contained no parsimony informative characters, were removed from further analyses. This resulted in 1,730 loci for the full analysis (see Table S7 for gene counts used in four-taxon tests).

IQ-TREE v2-rc1 was used with all 1,730 aligned loci to estimate a maximum likelihood concatenated (ML-CONCAT) tree with an edge-linked, proportional-partition model and 1,000 ultrafast bootstrap replicates [39,123,124]. The full IQ-TREE commandline used was: “iqtree -p Directory_of_Gene_Alignments --prefix -m MFP -c 8 - B 1000”. Maximum likelihood gene trees were estimated for each alignment with nucleotide substitution models selected using ModelFinder [125] as implemented in IQ-TREE. The full IQ-TREE commandline used was: “iqtree -s Directory_of_Gene_Alignments --prefix -m MFP -c 8”. We used the resulting maximum likelihood gene trees to estimate a species tree using ASTRAL III (ML-ASTRAL) [38]. Parsimony gene trees were generated using MPboot [77] and used to estimate a species tree using ASTRAL III (MP-ASTRAL), while PAUP* [74] was used to estimate the concatenated parsimony tree (MP-CONCAT) with 500 bootstrap replicates. IQ-TREE was used to calculate both gene concordance factors (gCFs) and site concordance factors (sCFs), with sCFs estimated from 300 randomly sampled quartets using the commandline: “iqtree --cf-verbose --gcf 1730_GENETREE.treefile -t Species_tree_file --df-tree --scf 300 -p Directory_of_Gene_Alignments -c 4”.

### Introgression Analyses

For each internal branch of the Primate tree where the proportion of discordant trees was > 5% of the total, concordance factors were used to calculate the test statistic Δ, where:

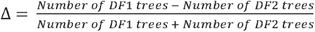

Where *DF1* trees represent the most frequent discordant topology and *DF2* trees are the second most frequent discordant topology. This is a normalized version of the statistic proposed by Huson *et al*. [36], which only included the numerator of this expression. Note also that, by definition, Δ here is always equal to or greater than 0. To test whether deviations from zero were significant (i.e. Δ > 0), we calculated Δ for 2,000 pseudo-replicate datasets generated by resampling gene trees with replacement. The resulting distribution was used to calculate *Z*-scores and the resulting *P*-values for the observed Δ value associated with each branch tested [126]. Of the 17 internal branches where > 5% of topologies were discordant, 7 were significant at *P* < 0.05, and selected for more extensive testing. For each of the 7 significant branches in the all-Primates tree, 4 taxa were selected that included the target branch as an internal branch. Single-copy genes present in each taxon were aligned as previously described. Alignments with no variant or parsimony-informative sites were removed from the analysis and gene trees were estimated using maximum likelihood in IQ-TREE 2. The test statistic, Δ, was calculated and significance was again determined using 2,000 bootstrap replicates with the *P*-value threshold for significance corrected for multiple comparisons (*m*=17) using the Dunn–Šidák correction [127,128].

### Molecular Dating

Molecular dating analyses were performed on 10 datasets consisting of 40 CDS alignments each sampled randomly without replacement from the 1,730 loci used to estimate the species tree. Gene alignments were concatenated into 10 supermatrices ranging from 36.7 kb – 42.7 kb in length (see Table S4 for the length of each alignment). Each dataset was then analyzed using PhyloBayes 3.3 [81] with sequences modeled using a site-specific substitution process with global exchange rates estimated from the data (CAT-GTR; [129]). Among-site rate-variation was modeled using a discrete gamma distribution with six rate categories. A relaxed molecular clock [130] with eight, soft-bounded, fossil calibrations (see Table S5) was used to estimate divergence times on the fixed species tree topology (Figure 1), the analyses were executed using the following command line: pb -x 1 15000 -d Alignment.phy -T Tree_file.tre -r outgroup_file.txt -cal 8_fossil.calib -sb -gtr -cat -bd -dgam 6 -ln -rp 90 90. Each dataset was analyzed for 15,000 generations, sampling every 10 generations, with 5,000 generations discarded as burn-in. Each dataset was analyzed twice to ensure convergence of the average age estimated for each node (Figure 3 shows the node age for both runs). To determine the effect of including a maximum constraint on the root of the Primates, we analyzed each dataset a third time with this constraint removed. Both the constrained and unconstrained node ages for major groups within the Primates are reported in Table 2.

## Acknowledgements

We thank Yue Liu for assistance in assembling the genomes, and Fábio Mendes and Gregg Thomas for helpful advice. This work was supported by National Science Foundation grants DBI-1564611 and DEB-1936187 (M.W.H), a Chan-Zuckerberg Initiative grant for Essential Open Source Software for Science (B.Q.M. and R.L.), and Australian Research Council grant DP-200103151 (R.L., B.Q.M., and M.W.H.).

## Supplementary Information

**Table S1**. Genomes analyzed in this study with the original NCBI release date, the publication for the reference used, and the accession number for the assembly. When possible the most recent version for each genome was used.

**Table S2**. All published genomes used in this study, including links to the assemblies and NCBI BioProjects. Annotation information is included for each genome at the time of download.

**Table S3**. Gaps/Ambiguities by species, and as a percentage of total alignment length. * denotes species sequenced this study.

**Table S4**. Lengths for each 40-locus concatenated alignment used in the molecular dating analyses. Each dataset was analyzed twice until node age estimates converged (15-25k steps) using a log-normal auto-correlated model (Thorne et al. 1998).

**Table S5**. Fossil calibrations employed in this study. Node numbering corresponds to the numbering in Figure 3. Median underflow/overflow for each calibration was calculated from 20 independent runs performed on 10 datasets (2 runs per dataset).

**Table S6**. Mean node age for 20 independent Phylobayes dating runs. Node numbers correspond to the numbering in Figure 3. The 95% HPD intervals were calculated by averaging the minimum and maximum of the 95% HPD interval for each dating run.

**Table S7**. Quartets used to test for significant Δ values for internal branches of the primate tree. Branches tested correspond to the labeled branches in Figure 3. After correcting for multiple comparisons (Dunn-Šidák, *P* = 0.00301), three internal branches and 8 quartets were found to have significant Δ values, indicating a likely introgression event.

**Supplementary Figure S1.**
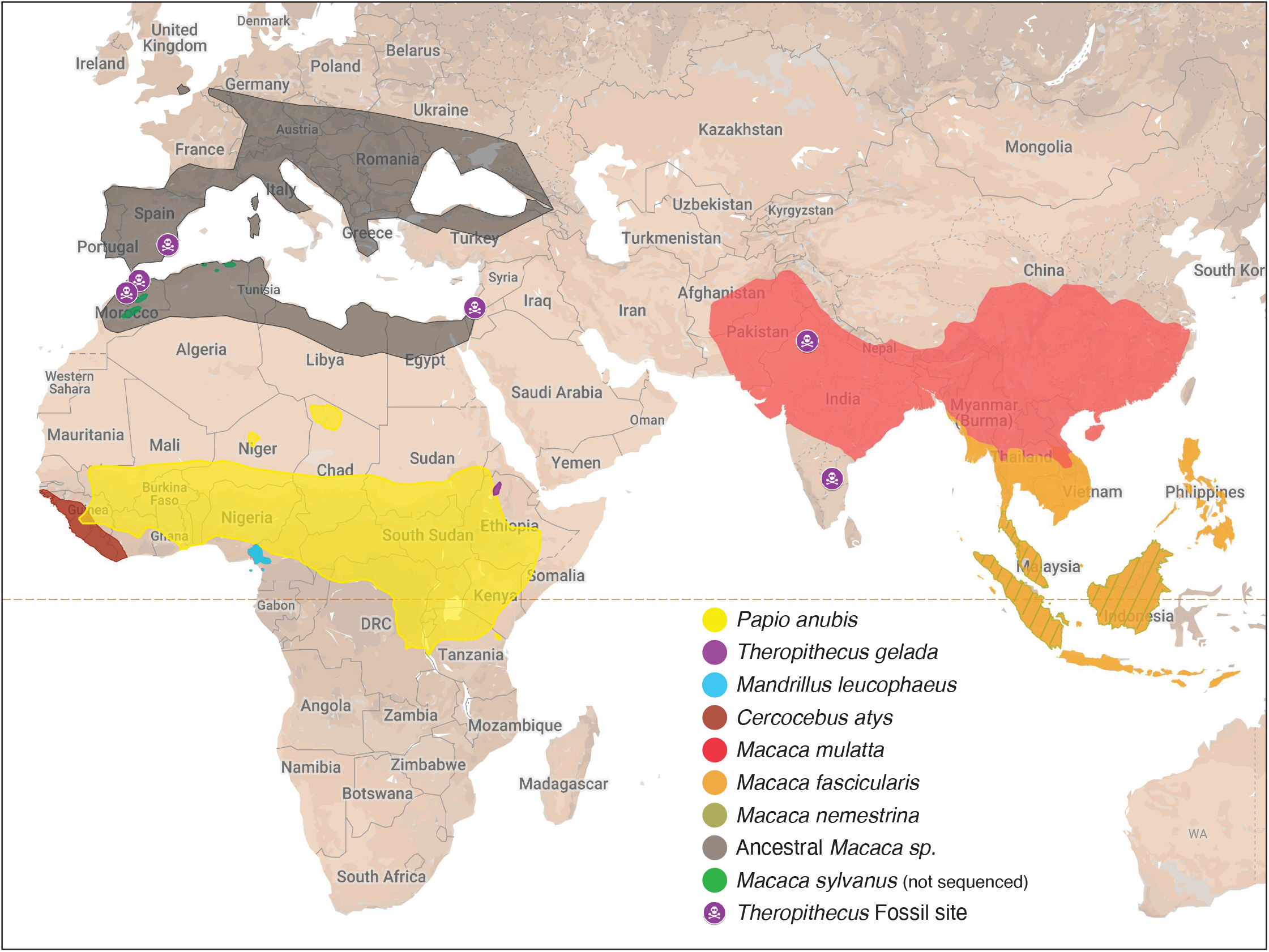
Present day species distributions for four African Papionini (*Papio, Theropithecus, Mandrillus*, and *Cercocebus*) and three Asian *Macaca* species included in the introgression analysis. The ancestral *Macaca* distribution (grey shading) is inferred from *Macaca* fossil localities in Africa and Europe as reviewed in Roos et al. (2019). The ancestral *Macaca* distribution likely represents only a fraction of the species range from the late Miocene to the late Pleistocene in Africa and Europe. The contemporary distribution of the African *Macaca sylvanus* (bright green) is included for reference. Fossil localities for *Theropithecus* species hypothesized to overlap contemporaneously with various ancestral *Macaca* are included. Citations for spatial data of extant species: *Macaca nemistrina* (Richardson et al., 2008), *Macaca fascicularis* (Ong & Richardson, 2008), *Macaca sylvanus* (Butynski et al., 2008), *Macaca mulatta* (Timmins et al., 2008), *Theropithecus gelada* (Gippoliti et al., 2019), *Papio anubis* (Kingdon et al., 2008), *Cercocebus atys* (Oates et al., 2016), and *Mandrillus leucophaeus* (Oates & Butynski, 2008).

