## Supplemental Tables 1, 3-7 for "Primate phylogenomics uncovers multiple rapid radiations and ancient interspecific introgression"

### Supplementary Materials

| Species | NCBI Year | Reference | Accession Used this Study |
|---|---|---|---|
| *Aotus nancymaae* (Ma's night monkey) | 2015 | (Thomas et al., 2018) | GCF_000952055.2 |
| *Callithrix jacchus* (White-tufted-ear marmoset) | 2010 | (Marmoset Genome Sequencing and Analysis Consortium, 2014) | GCF_000004665.1 |
| *Carlito syrichya* (Philippine tarsier) | 2013 | (Schmitz et al., 2016) | GCF_000164805.1 |
| *Cebus capucinus imitator* (White-faced sapajou) | 2016 | Dr. Amanda Melin, Washington University St. Louis;<br>Dr. Shoji Kawamura, University of Tokyo;<br>Dr. Wesley Warren, McDonnell Genome Institute;<br>Washington University School of Medicine, Unpublished | GCF_001604975.1 |
| *Cercocebus atys* (Sooty mangabey) | 2015 | (Palesch et al., 2018) | GCF_000955945.1 |
| *Chlorocebus sabaeus* (Green monkey) | 2014 | (Warren et al., 2015) | GCF_000409795.2 |
| ***Colobus angolensis palliatus* (Black and white colobus)** | **2015** | **This Study** | **GCF_000951035.1** |
| *Gorilla gorilla gorilla* (Western gorilla) | 2006 | (Scally et al., 2012) | GCF_000151905.2 |
| *Homo sapiens* | 2001 | (Church et al., 2011) | GCF_000001405.38 (GRCh38.p12) |
| *Macaca fascicularis* (Crab-eating macaque) | 2011 | (Yan et al., 2011) | GCF_000364345.1 |
| ***Macaca nemestrina* (Pig-tailed macaque)** | **2015** | **This Study** | **GCF_000956065.1** |
| *Macaca mullata* (Rhesus macaque) | 2006 | (Zimin et al., 2014) | GCF_000772875.2 |
| ***Mandrillus leucophaeus* (Drill)** | **2015** | **This Study** | **GCF_000951045.1** |
| *Microcebus murinus* (Gray mouse lemur) | 2007 | (Larsen et al., 2017) | GCF_000165445.2 |
| *Nomascus leucogenys* (Northern white-cheeked gibbon) | 2010 | (Carbone et al., 2014) | GCF_000146795.2 |
| *Otolemur garnetti* (Small-eared galago) | 2006 | Broad Institute, 2011, Unpublished | GCF_000181295.1 |
| *Pan paniscus* (Pygmy chimpanzee) | 2012 | (Prüfer et al., 2012) | GCF_000258655.2 |
| *Pan troglodytes* | 2005 | (Chimpanzee Sequencing and Analysis Consortium, 2005) | GCF_002880755.1 |
| *Papio anubis* (Olive baboon) | 2012 | (Rogers et al., 2019) | GCF_000264685.3 |
| *Piliocolobus tephrosceles* (Ugandan red colobus) | 2017 | University of Oregon, 2017, Unpublished | GCF_002776525.1 |
| *Pongo abelii* (Sumatran orangutan) | 2006 | University of Washington, 2017, Unpublished | GCF_002880775.1 |
| *Propithecus coquereli* (Coquerel's sifaka) | 2015 | Baylor College of Medicine, 2015, Unpublished | GCF_000956105.1 |
| *Rhinopithecus bieti* (Black snub-nosed monkey) | 2016 | (Yu et al., 2016) | GCF_001698545.1 |
| *Rhinopithecus roxellana* (Golden snub-nosed monkey) | 2014 | (Zhou et al., 2014) | GCF_000769185.1 |
| *Saimiri boliviensis boliviensis* (Bolivian squirrel monkey) | 2011 | Broad Institute, 2011, Unpublished | GCF_000235385.1 |
| *Theropithecus gelada* (Gelada) | 2018 | University of Washington, 2018, Unpublished | GCF_003255815.1 |
| *Tupaia chinensis* (Chinese tree shrew) | 2013 | (Fan et al., 2013) | GCF_000334495.1 |
| *Mus musculus* C57BL/6J (House Mouse) | 2002 | (Mouse Genome Sequencing Consortium et al., 2002) | GCF_000001635.26, GRCm38.p4 (Annotation release 106, 2016) |
| *Galeopterus variegatus* (Sunda flying lemur) | 2014 | Washington University, 2014, Unpublished | GCF_000696425.1 |

**Table S1**. Genomes analyzed in this study with the original NCBI release date, the publication for the reference used, and the accession number for the assembly. When possible the most recent version for each genome was used.

| Species | #Gap/Ambiguity | Percentage |
|---|---|---|
| *Macaca mulatta* | 71,090 | 4.04% |
| *Rhinopithecus roxellana* | 80,328 | 4.56% |
| *Chlorocebus sabaeus* | 88,366 | 5.02% |
| *Macaca nemestrina** | 88,650 | 5.03% |
| *Pongo abelii* | 89,290 | 5.07% |
| *Macaca fascicularis* | 97,942 | 5.56% |
| *Pan troglodytes* | 101,013 | 5.74% |
| *Cebus capucinus imitator* | 101,636 | 5.77% |
| *Cercocebus atys* | 102,521 | 5.82% |
| *Papio anubis* | 110,028 | 6.25% |
| *Aotus nancymaae* | 110,052 | 6.25% |
| *Microcebus murinus* | 114,128 | 6.48% |
| *Piliocolobus tephrosceles* | 115,512 | 6.56% |
| *Rhinopithecus bieti* | 119,118 | 6.76% |
| *Propithecus coquereli* | 120,633 | 6.85% |
| *Theropithecus gelada* | 128,355 | 7.29% |
| *Otolemur garnettii* | 128,999 | 7.33% |
| *Saimiri boliviensis* | 143,099 | 8.13% |
| *Gorilla gorilla* | 143,787 | 8.17% |
| *Callithrix jacchus* | 148,292 | 8.42% |
| *Pan paniscus* | 155,109 | 8.81% |
| *Homo sapiens* | 159,941 | 9.08% |
| *Colobus angolensis palliatus** | 177,747 | 10.09% |
| *Nomascus leucogenys* | 188,893 | 10.73% |
| *Galeopterus variegatus* | 199,749 | 11.34% |
| *Mus musculus* | 201,118 | 11.42% |
| *Tupaia chinensis* | 201,166 | 11.42% |
| *Mandrillus leucophaeus** | 204,583 | 11.62% |
| *Carlito syrichta* | 323,435 | 18.37% |

**Table S3**. Gaps/Ambiguities by species, and as a percentage of total alignment length.
* denotes species sequenced this study.

| Dataset | Length (bp) |
|---|---|
| Dataset 1 | 39,675 |
| Dataset 2 | 40,908 |
| Dataset 3 | 41,703 |
| Dataset 4 | 42,137 |
| Dataset 5 | 40,227 |
| Dataset 6 | 35,898 |
| Dataset 7 | 36,750 |
| Dataset 8 | 35,893 |
| Dataset 9 | 37,805 |
| Dataset 10 | 42,748 |

**Table S4**. Lengths for each 40-locus concatenated alignment used in the molecular dating analyses. Each dataset was analyzed twice until node age estimates converged (15-25k steps) using a log-normal auto-correlated model (Thorne et al. 1998).

| Node (Descendent Lineages) | Minimum Age MYA (citation) | Median Underflow (stdev) | Maximum Age (MYA) | Median Overflow (stdev) |
|---|---|---|---|---|
| Node 5 (Simiiformes, Tarsiiformes) | 43 (Franzen et al., 2009; Perelman et al., 2011; Poux & Douzery, 2004) | 0 (1.48) | NA | NA |
| Node 6 (Strepsirrhini, Haplorrhini) | 55.8 (Bloch et al., 2007; Sigé et al., 1990) | 0.87% (0.43) | 65.8 (Benton et al., 2015) | 3.37% (1.36) |
| Node 7 (Dermoptera, Primates) | 61<br>(Benton et al., 2015; Clemens & Wilson, 2009) | 5% (3.26) | 165 (Benton et al., 2015) | 0 (0.29) |
| Node 9 (Lorisiformes, Lemuriformes) | 38 (Seiffert et al., 2003) | 0.15% (0.65) | 56 (Benton et al., 2015; Sigé et al., 1990) | 0.19% (0.39) |
| Node 11 (Catarrhini) | 25 (Stevens et al., 2013) | 2% (1.48) | 34 (Benton et al., 2015) | 0.34% (2.58) |
| Node 13 (*Pongo, Homo*) | 14 (Raaum et al., 2005) | 0% (0.79) | 34 (Benton et al., 2015) | 0% |
| Node 15 (*Homo, Pan*) | 5.7 (Brunet et al., 2002) | 0.12% (0.62) | 10 (Benton et al., 2015) | 3.69% (2.58) |
| Node 22 (*Macaca, Papio*) | 7 (Steiper et al., 2004) | 1.28% (7.16) | NA | NA |

**Table S5**. Fossil calibrations employed in this study. Node numbering corresponds to the numbering in Figure 3. Median underflow/overflow for each calibration was calculated from 20 independent runs performed on 10 datasets (2 runs per dataset).

| Node Number | Mean Node Age (stdev) | Mean 95% Credibility Intervals |
|---|---|---|
| 1 | 16.97 (1.20) | 12.91 - 21.75 |
| 2 | 15.12 (1.17) | 11.25 - 19.59 |
| 3 | 17.21 (1.19) | 13.09 - 22.00 |
| 4 | 36.57 (0.59) | 32.16 - 41.41 |
| 5 | 59.01 (0.66) | 53.77 - 63.38 |
| 6 | 61.68 (0.53) | 56.23 - 65.90 |
| 7 | 66.57 (0.91) | 60.37 - 72.20 |
| 8 | 68.92 (1.69) | 62.10 - 75.75 |
| 9 | 47.74 (0.99) | 40.44 - 54.23 |
| 10 | 31.04 (2.02) | 20.95 - 40.86 |
| 11 | 28.41 (0.92) | 25.06 - 32.84 |
| 12 | 21.38 (0.74) | 17.96 - 25.47 |
| 13 | 19.19 (0.80) | 15.93 - 22.99 |
| 14 | 10.94 (0.73) | 8.48 - 13.15 |
| 15 | 8.54 (0.45) | 6.48 - 10.05 |
| 16 | 3.27 (0.82) | 2.18 - 4.54 |
| 17 | 16.79 (1.30) | 13.45 - 20.98 |
| 18 | 12.24 (1.32) | 9.16 - 16.17 |
| 19 | 10.14 (1.35) | 7.35 - 13.76 |
| 20 | 3.59 (0.95) | 2.22 - 5.77 |
| 21 | 12.83 (1.48) | 9.85 - 16.73 |
| 22 | 9.99 (1.57) | 7.46 - 13.44 |
| 23 | 5.65 (1.33) | 3.85 - 8.28 |
| 24 | 4.01 (1.28) | 2.61 - 6.06 |
| 25 | 8.53 (1.45) | 6.22 - 11.71 |
| 26 | 7.33 (1.49) | 5.21 - 10.22 |
| 27 | 5.85 (1.06) | 4.05 - 8.40 |

**Table S6**. Mean node age for 20 independent Phylobayes dating runs. Node numbers correspond to the numbering in Figure 3. The 95% HPD intervals were calculated by averaging the minimum and maximum of the 95% HPD interval for each dating run.

| $(((P_1,P_2),P_3),O)$ (Branch #, Figure 3) | Gene Trees | DF1/DF2 Counts, *P*-value | Significant at Dunn-Šidák *P* = 0.00301 |
|---|---|---|---|
| (((*Carlito, M. mulatta*), *Otolemur*),*Galeopterus*) (# 5) | 4,908 | 1170/1150, *p* = 0.687 | ☒ |
| (((*Homo, Pongo*), *Nomascus*), *M. mulatta*) (# 13) | 4,326 | 913/823, *p* = 0.030 | ☒ |
| (((*Homo, Chimp*), *Gorilla*), *Nomascus*) (# 15) | 4,099 | 703/635, *p* = 0.0625 | ☒ |
| (((*Chlorocebus, Papio*), *Colobus*), *Homo*) (# 51) | 6,358 | 709/646, *p* = 0.0854 | ☒ |
| (((*M. mulatta, M. nemistrina*),*Theropithecus*), *Colobus*) (# 23) | 5,109 | 216/194, *p* = 0.2710 | ☒ |
| (((*M. mulatta, M. fascicularis*), *M. nemistrina*), *Colobus*) (# 24) | 3,579 | 708/496, *p* = 1.373e -09 | ✓ |
| (((*Papio, Cercocebus*), *M. nemistrina*), *Colobus*) (# 25) | 4,290 | 863/821, *p* = 0.2998 | ☒ |
| (((*Papio, Cercocebus*), *M. mulatta*), *Colobus*) | 4,278 | 838/809, *p* = 0.4726 | ☒ |
| (((*Papio, Cercocebus*), *M. fascicularis*), *Colobus*) | 4,288 | 867/808, *p* = 0.1428 | ☒ |
| (((*Papio, Mandrillus*), *M. nemistrina*), *Colobus*) | 4,386 | 1119/764, *p* = 2.220e -16 | ✓ |
| (((*Papio, Mandrillus*), *M. mulatta*), *Colobus*) | 4,360 | 1099/739, *p* = 0.0 | ✓ |
| (((*Papio, Mandrillus*), *M. fascicularis*), *Colobus*) | 4,375 | 1125/735, *p* = 0.0 | ✓ |
| (((*Theropithecus, Cercocebus*), *M. nemistrina*), *Colobus*) | 4,323 | 993/766, *p* = 6.679e -08 | ✓ |
| (((*Theropithecus, Cercocebus*), *M. mulatta*), *Colobus*) | 4,303 | 975/736, *p* = 4.958e -09 | ✓ |
| (((*Theropithecus, Cercocebus*), *M. fascicularis*), *Colobus*) | 4,305 | 1000/748, *p* = 3.020e -09 | ✓ |
| (((*Theropithecus, Mandrillus*), *M. nemistrina*), *Colobus*) | 4,306 | 888/806, *p* = 0.0440 | ☒ |
| (((*Theropithecus, Mandrillus*), *M. mulatta*), *Colobus*) | 4,277 | 872/759, *p* = 0.0059 | ☒ |
| (((*Theropithecus, Mandrillus*), *M. fascicularis*), *Colobus*) | 4,261 | 866/771, *p* = 0.01 | ☒ |
| (((*Papio, Theropithecus*), *Cercocebus*), *Colobus*) (# 27) | 5,605 | 912/569, *p* = 0.0 | ✓ |

**Table S7.** Quartets used to test for significant Δ values for internal branches of the primate tree. Branches tested correspond to the labeled branches in Figure 3. After correcting for multiple comparisons (Dunn-Šidák, *P* = 0.00301), three internal branches and 8 quartets were found to have significant Δ values, indicating a likely introgression event.

**Supplementary Figure S1**. Present day species distributions for four African Papionini (*Papio*, *Theropithecus*, *Mandrillus*, and *Cercocebus*) and three Asian *Macaca* species included in the introgression analysis. The ancestral *Macaca* distribution (grey shading) is inferred from *Macaca* fossil localities in Africa and Europe as reviewed in Roos et al. (2019). The ancestral *Macaca* distribution likely represents only a fraction of the species range from the late Miocene to the late Pleistocene in Africa and Europe. The contemporary distribution of the African *Macaca sylvanus* (bright green) is included for reference. Fossil localities for *Theropithecus* species hypothesized to overlap contemporaneously with various ancestral *Macaca* are included. Citations for spatial data of extant species: *Macaca nemistrina* (Richardson et al., 2008), *Macaca fascicularis* (Ong & Richardson, 2008), *Macaca sylvanus* (Butynski et al., 2008), *Macaca mulatta* (Timmins et al., 2008), *Theropithecus gelada* (Gippoliti et al., 2019), *Papio anubis* (Kingdon et al., 2008), *Cercocebus atys* (Oates et al., 2016), and *Mandrillus leucophaeus* (Oates & Butynski, 2008).

### Supplementary Material References


Benton, M. J., Donoghue, P. C. J., Asher, R. J., Friedman, M., Near, T. J., & Vinther, J. (2015). Constraints on the timescale of animal evolutionary history. *Palaeontologia Electronica*, *18*(1.FC), 1–106. https://doi.org/10.26879/424

Bloch, J. I., Silcox, M. T., Boyer, D. M., & Sargis, E. J. (2007). New Paleocene skeletons and the relationship of plesiadapiforms to crown-clade primates. *Proceedings of the National Academy of Sciences*, *104*(4), 1159–1164. https://doi.org/10.1073/pnas.0610579104

Brunet, M., Guy, F., Pilbeam, D., Mackaye, H. T., Likius, A., Ahounta, D., Beauvilain, A., Blondel, C., Bocherens, H., Boisserie, J.-R., De Bonis, L., Coppens, Y., Dejax, J., Denys, C., Duringer, P., Eisenmann, V., Fanone, G., Fronty, P., Geraads, D., … Zollikofer, C. (2002). A new hominid from the Upper Miocene of Chad, Central Africa. *Nature*, *418*(6894), 145–151. https://doi.org/10.1038/nature00879

Butynski, T. M., Cortes, J., Waters, S., Fa, J., Hobbelink, M. E., van Lavieren, E., Belbachir, F., Cuzin, F., de Smet, K., Mouna, M., de Longh, H., Menard, N., & Camperio-Ciani, A. (2008). *Macaca sylvanus. The IUCN Red List of Threatened Species 2008: E.T12561A3359140.* https://dx.doi.org/10.2305/IUCN.UK.2008.RLTS.T12561A3359140.en.

Carbone, L., Harris, R. A., Gnerre, S., Veeramah, K. R., Lorente-Galdos, B., Huddleston, J., Meyer, T. J., Herrero, J., Roos, C., Aken, B., Anaclerio, F., Archidiacono, N., Baker, C., Barrell, D., Batzer, M. A., Beal, K., Blancher, A., Bohrson, C. L., Brameier, M., … Gibbs, R. A. (2014). Gibbon genome and the fast karyotype evolution of small apes. *Nature*, *513*(7517), 195–201. https://doi.org/10.1038/nature13679

Chimpanzee Sequencing and Analysis Consortium. (2005). Initial sequence of the chimpanzee genome and comparison with the human genome. *Nature*, *437*(7055), 69–87. https://doi.org/10.1038/nature04072

Church, D. M., Schneider, V. A., Graves, T., Auger, K., Cunningham, F., Bouk, N., Chen, H.-C., Agarwala, R., McLaren, W. M., Ritchie, G. R. S., Albracht, D., Kremitzki, M., Rock, S., Kotkiewicz, H., Kremitzki, C., Wollam, A., Trani, L., Fulton, L., Fulton, R., … Hubbard, T. (2011). Modernizing reference genome assemblies. *PLoS Biology*, *9*(7), e1001091. https://doi.org/10.1371/journal.pbio.1001091

Clemens, W. A., & Wilson, G. P. (2009). Early Torrejonian mammalian local faunas from northeastern Montana, USA. *Museum of Northern Arizona Bulletin*, *65*, 111–158.

Fan, Y., Huang, Z.-Y., Cao, C.-C., Chen, C.-S., Chen, Y.-X., Fan, D.-D., He, J., Hou, H.-L., Hu, L., Hu, X.-T., Jiang, X.-T., Lai, R., Lang, Y.-S., Liang, B., Liao, S.-G., Mu, D., Ma, Y.-Y., Niu, Y.-Y., Sun, X.-Q., … Yao, Y.-G. (2013). Genome of the Chinese tree shrew. *Nature Communications*, *4*, 1426. https://doi.org/10.1038/ncomms2416

Franzen, J. L., Gingerich, P. D., Habersetzer, J., Hurum, J. H., von Koenigswald, W., & Smith, B. H. (2009). Complete primate skeleton from the Middle Eocene of Messel in Germany: Morphology and paleobiology. *PloS One*, *4*(5), e5723. https://doi.org/10.1371/journal.pone.0005723

Gippoliti, S., Mekonnen, A., Burke, R., Nguyen, N., & Fashing, P. J. (2019). *Theropithecus gelada. The IUCN Red List of Threatened Species 2019: E.T21744A17941908.* https://dx.doi.org/10.2305/IUCN.UK.2019-3.RLTS.T21744A17941908.en.

Kingdon, J., Butynski, T. M., & De Jong, Y. (2008). *Papio anubis. The IUCN Red List of Threatened Species 2008: E.T40647A10348950.* https://dx.doi.org/10.2305/IUCN.UK.2008.RLTS.T40647A10348950.en.

Larsen, P. A., Harris, R. A., Liu, Y., Murali, S. C., Campbell, C. R., Brown, A. D., Sullivan, B. A., Shelton, J., Brown, S. J., Raveendran, M., Dudchenko, O., Machol, I., Durand, N. C., Shamim, M. S., Aiden, E. L., Muzny, D. M., Gibbs, R. A., Yoder, A. D., Rogers, J., & Worley, K. C. (2017). Hybrid de novo genome assembly and centromere characterization of the gray mouse lemur (Microcebus murinus). *BMC Biology*, *15*(1), 110. https://doi.org/10.1186/s12915-017-0439-6

Marmoset Genome Sequencing and Analysis Consortium. (2014). The common marmoset genome provides insight into primate biology and evolution. *Nature Genetics*, *46*(8), 850–857. https://doi.org/10.1038/ng.3042

Mouse Genome Sequencing Consortium, Waterston, R. H., Lindblad-Toh, K., Birney, E., Rogers, J., Abril, J. F., Agarwal, P., Agarwala, R., Ainscough, R., Alexandersson, M., An, P., Antonarakis, S. E., Attwood, J., Baertsch, R., Bailey, J., Barlow, K., Beck, S., Berry, E., Birren, B., … Lander, E. S. (2002). Initial sequencing and comparative analysis of the mouse genome. *Nature*, *420*(6915), 520–562. https://doi.org/10.1038/nature01262

Oates, J. F., & Butynski, T. M. (2008). *Mandrillus leucophaeus ssp. Leucophaeus. The IUCN Red List of Threatened Species 2008: E.T12756A3378112.* https://dx.doi.org/10.2305/IUCN.UK.2008.RLTS.T12756A3378112.en.

Oates, J. F., Gippoliti, S., & Groves, C. P. (2016). *Cercocebus atys. The IUCN Red List of Threatened Species 2016: E.T136933A92247942.* https://dx.doi.org/10.2305/IUCN.UK.2016-1.RLTS.T136933A92247942.en.

Ong, P., & Richardson, M. (2008). *Macaca fascicularis. The IUCN Red List of Threatened Species 2008: E.T12551A3355536.* https://dx.doi.org/10.2305/IUCN.UK.2008.RLTS.T12551A3355536.en.

Palesch, D., Bosinger, S. E., Tharp, G. K., Vanderford, T. H., Paiardini, M., Chahroudi, A., Johnson, Z. P., Kirchhoff, F., Hahn, B. H., Norgren, R. B., Patel, N. B., Sodora, D. L., Dawoud, R. A., Stewart, C.-B., Seepo, S. M., Harris, R. A., Liu, Y., Raveendran, M., Han, Y., … Silvestri, G. (2018). Sooty mangabey genome sequence provides insight into AIDS resistance in a natural SIV host. *Nature*, *553*(7686), 77–81. https://doi.org/10.1038/nature25140

Perelman, P., Johnson, W. E., Roos, C., Seuánez, H. N., Horvath, J. E., Moreira, M. A. M., Kessing, B., Pontius, J., Roelke, M., Rumpler, Y., Schneider, M. P. C., Silva, A., O'Brien, S. J., & Pecon-Slattery, J. (2011). A molecular phylogeny of living primates. *PLoS Genetics*, *7*(3), e1001342. https://doi.org/10.1371/journal.pgen.1001342

Poux, C., & Douzery, E. J. P. (2004). Primate phylogeny, evolutionary rate variations, and divergence times: A contribution from the nuclear gene IRBP. *American Journal of Physical Anthropology*, *124*(1), 1–16. https://doi.org/10.1002/ajpa.10322

Prüfer, K., Munch, K., Hellmann, I., Akagi, K., Miller, J. R., Walenz, B., Koren, S., Sutton, G., Kodira, C., Winer, R., Knight, J. R., Mullikin, J. C., Meader, S. J., Ponting, C. P., Lunter, G., Higashino, S., Hobolth, A., Dutheil, J., Karakoç, E., … Pääbo, S. (2012). The bonobo genome compared with the chimpanzee and human genomes. *Nature*, *486*(7404), 527–531. https://doi.org/10.1038/nature11128

Raaum, R. L., Sterner, K. N., Noviello, C. M., Stewart, C.-B., & Disotell, T. R. (2005). Catarrhine primate divergence dates estimated from complete mitochondrial genomes: Concordance with fossil and nuclear

DNA evidence. *Journal of Human Evolution*, *48*(3), 237–257. https://doi.org/10.1016/j.jhevol.2004.11.007

Richardson, M., Mittermeier, R. A., Rylands, A. B., & Konstant, B. (2008). *Macaca nemestrina. The IUCN Red List of Threatened Species 2008: E.T12555A3356892.* https://dx.doi.org/10.2305/IUCN.UK.2008.RLTS.T12555A3356892.en.

Rogers, J., Raveendran, M., Harris, R. A., Mailund, T., Leppälä, K., Athanasiadis, G., Schierup, M. H., Cheng, J., Munch, K., Walker, J. A., Konkel, M. K., Jordan, V., Steely, C. J., Beckstrom, T. O., Bergey, C., Burrell, A., Schrempf, D., Noll, A., Kothe, M., … Baboon Genome Analysis Consortium. (2019). The comparative genomics and complex population history of Papio baboons. *Science Advances*, *5*(1), eaau6947. https://doi.org/10.1126/sciadv.aau6947

Scally, A., Dutheil, J. Y., Hillier, L. W., Jordan, G. E., Goodhead, I., Herrero, J., Hobolth, A., Lappalainen, T., Mailund, T., Marques-Bonet, T., McCarthy, S., Montgomery, S. H., Schwalie, P. C., Tang, Y. A., Ward, M. C., Xue, Y., Yngvadottir, B., Alkan, C., Andersen, L. N., … Durbin, R. (2012). Insights into hominid evolution from the gorilla genome sequence. *Nature*, *483*(7388), 169. https://doi.org/10.1038/nature10842

Schmitz, J., Noll, A., Raabe, C. A., Churakov, G., Voss, R., Kiefmann, M., Rozhdestvensky, T., Brosius, J., Baertsch, R., Clawson, H., Roos, C., Zimin, A., Minx, P., Montague, M. J., Wilson, R. K., & Warren, W. C. (2016). Genome sequence of the basal haplorrhine primate Tarsius syrichta reveals unusual insertions. *Nature Communications*, *7*, 12997. https://doi.org/10.1038/ncomms12997

Seiffert, E. R., Simons, E. L., & Attia, Y. (2003). Fossil evidence for an ancient divergence of lorises and galagos. *Nature*, *422*(6930), 421–424. https://doi.org/10.1038/nature01489

Sigé, B., Jaeger, J.-J., Sudre, J., & Vianey-Liaud, M. (1990). Altiatlasius koulchii n. Gen. Et sp., primate omomyidé du Paléocène supérieur du Maroc, et les origines des euprimates. *Palaeontographica Abteilung A*, 31–56.

Steiper, M. E., Young, N. M., & Sukarna, T. Y. (2004). Genomic data support the hominoid slowdown and an Early Oligocene estimate for the hominoid–cercopithecoid divergence. *Proceedings of the National Academy of Sciences*, *101*(49), 17021–17026. https://doi.org/10.1073/pnas.0407270101

Stevens, N. J., Seiffert, E. R., O'Connor, P. M., Roberts, E. M., Schmitz, M. D., Krause, C., Gorscak, E., Ngasala, S., Hieronymus, T. L., & Temu, J. (2013). Palaeontological evidence for an Oligocene divergence between Old World monkeys and apes. *Nature*, *497*(7451), 611–614. https://doi.org/10.1038/nature12161

Thomas, G. W. C., Wang, R. J., Puri, A., Harris, R. A., Raveendran, M., Hughes, D. S. T., Murali, S. C., Williams, L. E., Doddapaneni, H., Muzny, D. M., Gibbs, R. A., Abee, C. R., Galinski, M. R., Worley, K. C., Rogers, J., Radivojac, P., & Hahn, M. W. (2018). Reproductive longevity predicts mutation rates in primates. *Current Biology : CB*, *28*(19), 3193–3197.e5. https://doi.org/10.1016/j.cub.2018.08.050

Timmins, R. J., Richardson, M., Chhangani, A., & Yongcheng, L. (2008). *Macaca mulatta. The IUCN Red List of Threatened Species 2008: E.T12554A3356486.* https://dx.doi.org/10.2305/IUCN.UK.2008.RLTS.T12554A3356486.en.

Warren, W. C., Jasinska, A. J., García-Pérez, R., Svardal, H., Tomlinson, C., Rocchi, M., Archidiacono, N., Capozzi, O., Minx, P., Montague, M. J., Kyung, K., Hillier, L. W., Kremitzki, M., Graves, T., Chiang, C., Hughes, J., Tran, N., Huang, Y., Ramensky, V., … Freimer, N. B. (2015). The genome of the vervet (Chlorocebus aethiops sabaeus). *Genome Research*, *25*(12), 1921–1933. https://doi.org/10.1101/gr.192922.115

Yan, G., Zhang, G., Fang, X., Zhang, Y., Li, C., Ling, F., Cooper, D. N., Li, Q., Li, Y., van Gool, A. J., Du, H., Chen, J., Chen, R., Zhang, P., Huang, Z., Thompson, J. R., Meng, Y., Bai, Y., Wang, J., … Wang, J.

(2011). Genome sequencing and comparison of two nonhuman primate animal models, the cynomolgus and Chinese rhesus macaques. *Nature Biotechnology*, *29*(11), 1019–1023. https://doi.org/10.1038/nbt.1992

Yu, L., Wang, G.-D., Ruan, J., Chen, Y.-B., Yang, C.-P., Cao, X., Wu, H., Liu, Y.-H., Du, Z.-L., Wang, X.-P., Yang, J., Cheng, S.-C., Zhong, L., Wang, L., Wang, X., Hu, J.-Y., Fang, L., Bai, B., Wang, K.-L., … Zhang, Y.-P. (2016). Genomic analysis of snub-nosed monkeys ( Rhinopithecus ) identifies genes and processes related to high-altitude adaptation. *Nature Genetics*, *48*(8), 947–952. https://doi.org/10.1038/ng.3615

Zhou, X., Wang, B., Pan, Q., Zhang, J., Kumar, S., Sun, X., Liu, Z., Pan, H., Lin, Y., Liu, G., Zhan, W., Li, M., Ren, B., Ma, X., Ruan, H., Cheng, C., Wang, D., Shi, F., Hui, Y., … Li, M. (2014). Whole-genome sequencing of the snub-nosed monkey provides insights into folivory and evolutionary history. *Nature Genetics*, *46*(12), 1303–1310. https://doi.org/10.1038/ng.3137

Zimin, A. V., Cornish, A. S., Maudhoo, M. D., Gibbs, R. M., Zhang, X., Pandey, S., Meehan, D. T., Wipfler, K., Bosinger, S. E., Johnson, Z. P., Tharp, G. K., Marçais, G., Roberts, M., Ferguson, B., Fox, H. S., Treangen, T., Salzberg, S. L., Yorke, J. A., & Norgren, R. B. (2014). A new rhesus macaque assembly and annotation for next-generation sequencing analyses. *Biology Direct*, *9*(1), 20. https://doi.org/10.1186/1745-6150-9-20
